# Stable Synthetic Organelles from Aqueous Two-Phase Systems with Access to the Cell Translation Machinery

**DOI:** 10.1101/2025.08.20.671219

**Authors:** Marcos Masukawa, Lea Duttenhofer, Soumya Sethi, Christoph Drees, Andreas Walther

**Author notes:** equal contribution (Marcos Masukawa, Lea Duttenhofer).

## Abstract

Cells compartmentalize vital processes in membrane-less organelles to gain spatiotemporal control of metabolism, signaling, and for protection under stress. While such compartments can be manipulated or even *de novo* designed with genetic engineering of cells, the transfer and operation of exogenous synthetic compartments for intracellular engineering is challenged by developing pathways for implantation into the cell, stability issues, toxicity, ability to maintain compartmentalization, and access to cellular machinery. Here we introduce dextran-lipid droplets as versatile exogenous synthetic compartments, that are readily uptaken by model cancer and immune cells and are stable inside cells for days. Furthermore, the droplets can encapsulate nucleic acids with high efficiency, are non-toxic and endosomal escape occurs when formulated with ionizable lipids, as shown by reporter protein translation of an mRNA hosted inside the organelles. We propose that such synthetic microreactors implanted into the cells will become an important bioengineering tool to incorporate more complex and bioorthogonal molecular systems into cells.

## Introduction

Cells possess membraneless organelles that are formed by liquid-liquid phase separation (LLPS) and consist of proteins^[1]^, RNA^[1]^, and lipids.^[2]^ Such compartments act as hubs for segmenting and processing information, such as for metabolites, catalytic reactions, and signaling molecules.^[1]^ Designer proteins that undergo phase separation can generate analogous structures that recruit and release molecules to alter cell behavior.^[1,3]^ The ability to alter cell physiology through the formation of new synthetic compartments is one of many challenges of intracellular matrix engineering, aiming to tune the properties of the cell interior by creating new structures and altering signaling or metabolism. In recent years, new tools have been emerging to control subcellular components due to advances, for instance, in soft matter and DNA nanotechnology.^[4]^ An ongoing effort of the field is the creation of functional synthetic compartments, that is, synthetic organelles.^[4,6,7]^

There are two main approaches to the creation of synthetic organelles:^[5]^ On the one hand, genetic engineering of cells can produce proteins that undergo LLPS, allowing for the *de novo* fabrication of functional compartments.^[3,6]^ This approach is becoming increasingly robust but may ultimately face some limitations for applications due to genetic engineering, as well as it does not permit to install bioorthogonal synthetic functionalities. On the other hand, synthetic compartments can be formed outside cells and transferred into them. Here, polymer capsules,^[7]^ polymersomes,^[8]^ and protein cages^[9]^ present viable entry points for installing synthetic compartments in eukaryotic cells.^[5]^ For example, polymersomes, when appropriately functionalized with cell-penetrating TAT^[8]^ or RGD^[10]^ peptides, can be localized into cells. One application is the use of polymersomes to increase the resistance of HEK-293 cells to oxidative damage by loading the polymersomes with catalase.^[8]^ Nevertheless, such membrane-bound polymersomes, capsules, and protein cages differ considerably from membrane-less condensate organelles of cells with respect to stability, permeability, and other properties such as molecularly crowded condensates.

Here we introduce exogenous, non-crosslinked dextran-lipid condensate compartments – of high analogy to cellular membraneless organelles although not being composed of proteins – which are efficiently uptaken by eukaryotic cells and lead to intracellular synthetic organelles that are stable for days. During this period, the droplets do not fuse, and the cells do not secrete them. During preparation, such droplets spontaneously take up certain biomolecules, such as mRNA. Dextran-lipid droplets prepared from ionizable lipids allow for protein translation from the mRNA cargo, confirming that these structures are not trapped in endosomes but have access to the translation machinery. The mRNA remains compartmentalized during this process. Next to investigations in HeLa cancer cells, we also demonstrate successful uptake and protein translation for the model immune cells Jurkat (T-cell model) and THP-1 (macrophage model). Together, this strategy presents a straightforward method for creating a non-proteinaceous condensate in cells, and reinforces the potential of self-assembling coacervates in intracelullar engineering.^[11,12]^ We believe that orthogonality, stability, variability, and facile integration into the cytosol underpin the future use of dextran-lipid droplets as synthetic organelles.

## Results and Discussion

### Formation of Dextran-Lipid Droplets Assisted by Aqueous Two-Phase Systems

We use dextran as the basis for fabricating synthetic compartments to be transferred into cells. The motivation to use dextran builds upon its ease of chemical functionalization and proven safety at the *in vivo* level^[13]^ up to clinical trials.^[14]^ Dextran has low immunogenicity and its appropriate lipophilic/hydrophilic balance enables loading with various drugs and (bio)macromolecules.^[15]^ Additionally, various cancer cell lines^[16]^ and immune cells^[17]^ uptake molecular dextran as mediated by class A scavenger receptors or carbohydrate receptors such as mannose receptors.^[18,19]^ In general, cells uptake molecular dextran with a molecular weight up to ∼150 kDa^[20]^ and direct the dextran to an endolysosomal pathway.^[21,22]^ This means that unmodified dextrans do not integrate much into the cytosol. Dextrans of very large molecular weights (e.g. > 500 kDa) have been introduced into the cell cytoplasm by microinjection^[23]^ or by modifying the dextran with thiol groups, leading to thiol mediated uptake.^[19]^ Once inside the cell, large dextrans are stable over several days but do not form compartments.^[23,24]^ Modified dextrans have been introduced in cells as crosslinked particles and are often engineered for degradability,^[25–28]^ functioning rather as degradable nanocarriers but not as dynamic intracellular compartments.

Our approach for building nano/micro-scale dextran droplets uses an aqueous two-phase systems (ATPS) formed by spontaneous demixing of dextran/polyethylene glycol (DEX/PEG) solutions.^[29–31]^ In more details, we prepared ATPSs of PEG (6 kDa, 100 g/L) and DEX (450-600 kDa, 71 µg/mL), leading to micrometer-sized DEX droplets after shaking. We used spectrally distinct dye-labeled dextrans, DEX_FITC_ and DEX_RITC_ (at 8 µg/mL), for visualizing these droplets, tracking different droplet populations, and excluding dye effects. One of the major benefits of DEX/PEG ATPS is the ease of incorporating biomolecules,^[32]^ for which we used – for specific samples – a mRNA encoding eGFP (8 µg/mL), a fluorescently labeled mRNA_AZ405-rUTP_ also encoding eGFP (8 µg/mL) and a mRNA encoding mCherry (16 µg/mL). A small quantity of diethylaminomethyl(DEAE)-DEX (0.8 µg/mL) is present to minimize mRNA degradation^[33]^ and enhance compartmentalization in all formulations. Tables **S1-4** give an overview of all components and concentrations used in the experiments.

To prevent coalescence of the DEX-PEG ATPS, we stabilized the interface by lipids^[34]^ through a lipid-film rehydration method (details in methods). Previous studies showed that such lipid-stabilized ATPSs do not have a defined structure such as a bilayer, but still efficiently stabilize interfaces.^[31]^ We used two different lipid formulations: (1) 1,2-Dioleoyl-sn-glycero-3-phosphocholine (DOPC) and (2) an ionizable lipid mixture (ILM, composition in methods) used for the formation of lipid nanoparticles for mRNA vaccines and aiding in endosomal escape after cellular uptake.^[35]^ When necessary, we added trace amounts of two dye-labeled lipids (DOPE_Atto488_ and PE_Rhod_; abbreviations in Tables **S1**,**2**). After emulsification of the lipid film with the ATPS, we diluted such lipid-stabilized ATPS 1:1 in cell culture medium and extruded them through polycarbonate membranes with pore sizes of 50, 400 or 1000 nm to reduce the size of the DEX droplets (Figure **1a**). After extrusion, we refer to lipid-stabilized ATPS as DEX-lipid droplets. The droplets are further diluted 1:1 in cell culture medium, which results in a final concentration of approximately 90 µg/mL of DEX-lipid droplets.

**Figure 1.**
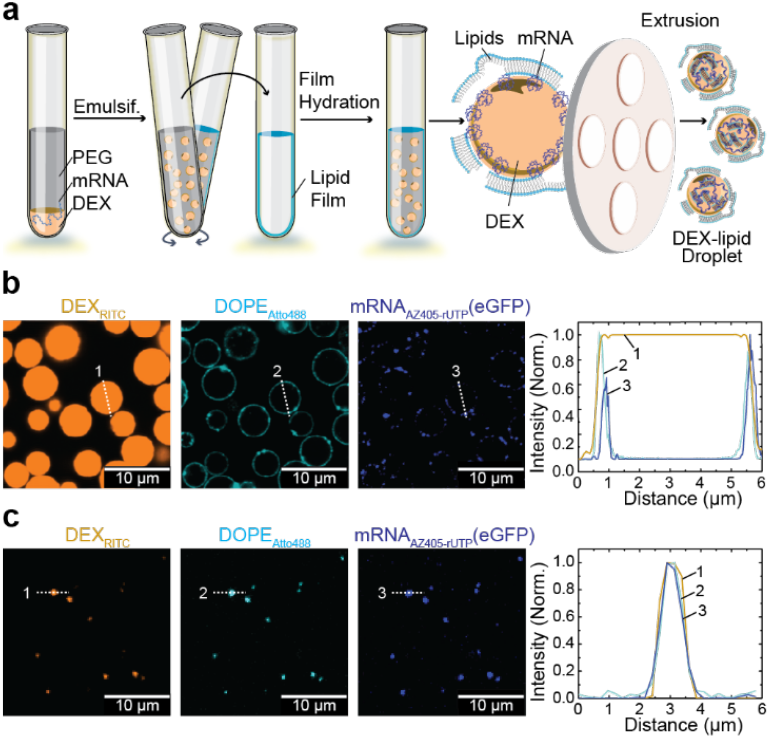
Preparation of DEX-lipid droplets with mRNA cargo from DEX-PEG ATPS. **(a)** Emulsification of DEX, PEG, and mRNA and addition to a lipid film. Extrusion of the lipid-stabilized DEX droplets reduces the droplets to the desired size. The DEX-lipid droplets become stable against dilution. **(b)** CLSM images of lipid-stabilized ATPS (no extrusion) of DEX/DEX_RITC_/mRNA_AZ405-rUTP_-ILM/DOPE_Atto448_, showing intensity profile of each fluorescent marker. **(c)** CLSM images of DEX-lipid droplets of DEX/DEX_RITC_/mRNA_AZ405-rUTP_-ILM/DOPE_Atto448_ extruded with through a 1000 nm membrane, showing intensity profile of each fluorescent marker. See Tables **S3**,**4** for compositions of samples in **b** and **c**.

Lipid-stabilized ATPS and the extruded DEX-lipid droplets have different structures as seen by Confocal Laser Scanning Microscopy (CLSM). The lipid-stabilized ATPS has an accumulation of lipids and mRNA at the interface (Figure **1b**). The lipid accumulation is qualitatively similar to previously reported lipid-stabilized ATPS.^[31]^ However, after extrusion, the corresponding DEX-lipid droplets display homogenous condensate structures, as for instance seen for a DEX/DEX_RITC_/mRNA_AZ405-rUTP_-ILM/DOPE_Atto448_ emulsion after extrusion through a 1000 nm membrane via the cross-sectional analysis (Figure **1c**). In addition, the membrane pore size used for extrusion controls the final DEX-lipid droplet size. Dynamic Light Scattering of DEX-ILM droplets yields z-average diameters of 50 ± 4 nm, 412 ± 31 nm, and 942 ± 119 nm when extruded through 50, 400, and 1000 nm membranes, respectively. This demonstrates a simple and robust control over the DEX droplet size.

### Cell Uptake Depends on DEX-Lipid Droplet Size

Next, we discuss cellular uptake of differently sized DEX-lipid droplets for eukaryotic cells, where we selected HeLa cells as robust model cell line. In a typical experiment, we seeded 2×10^3^ cells in 8-well slides and added 300 µL of DEX-lipid droplets (90 µg/mL). Quantification of CLSM data and flow cytometry shows that the uptake of DEX-lipid droplets prepared either with DOPC or ILM is size dependent (Figure **2**). For DEX-ILM, the average uptake measured by fluorescence intensity (FI) is positively correlated with droplet size. The difference between DEX-ILM droplets prepared by extrusion through 50 and 400 nm pores is not significant, while the uptake of those prepared by extrusion through 1000 nm is significantly higher. Size and uptake also correlate for DEX-DOPC droplets. In this case, the difference between droplets prepared through extrusion pores of 50 and 400 nm is already significant, but not between 400 and 1000 nm pores (Figures **2a-c** for DEX/DEX_RITC_-ILM, Figure **S1** for DEX-DOPC/PE_Rhod_ and DEX/DEX_RITC_-DOPC, see Table **S5** for significance definition). The positive correlation between size suggests macropinocytosis as the predominant uptake mechanism.^[36,37]^ This is in line with the observation that high molecular weight dextrans (>70 kDa) are known to be uptaken almost entirely by macropinocytosis.^[38]^ Live cell imaging of the uptake process depicts droplets adhering to extended cell structures, such as filopodia, that are transported towards the cell (see Figure **2d**, Supplementary Video **1-3**). Transport along filipodia is one of the main uptake pathways of exosomes^[39,40]^ and viruses,^[41]^ but seldomly reported for polymeric particles.^[42]^

**Figure 2.**
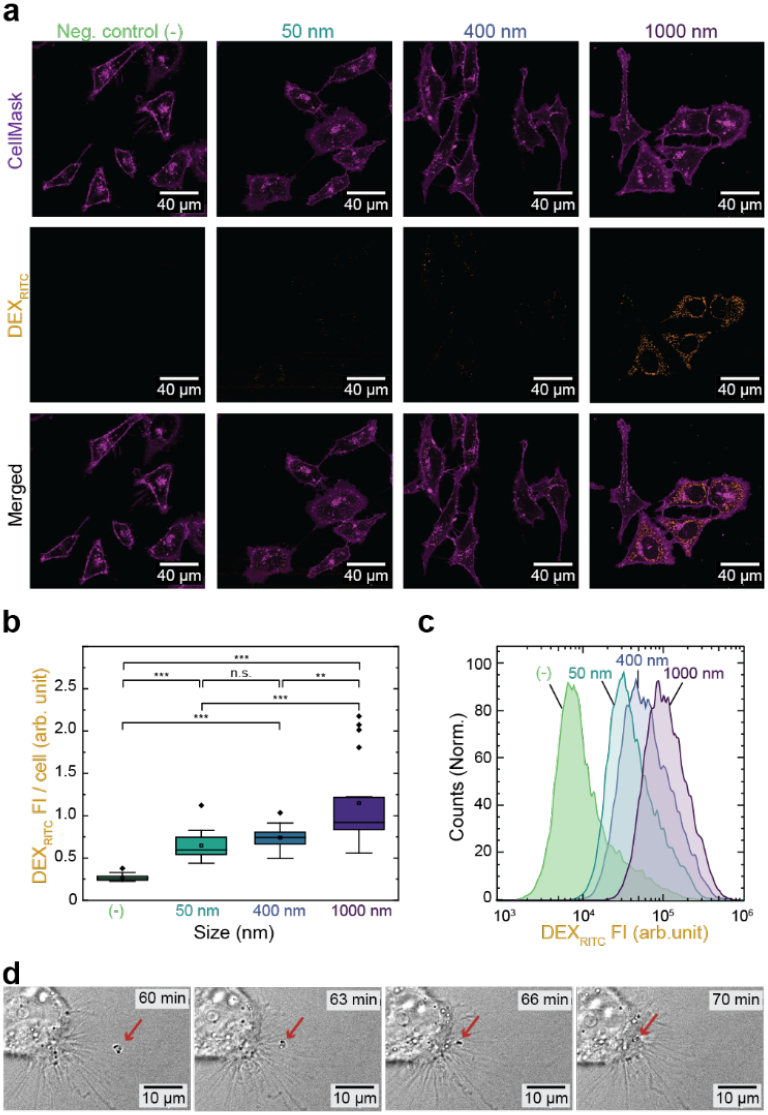
Uptake of DEX-ILM droplets by HeLa cells according to extrusion size. **(a)** CLSM images of cells 24 h after treatment with DEX/DEX_RITC_-ILM. Cells are stained at the membrane using CellMask. Culture medium added as negative control. **(b)** DEX_RITC_ fluorescence intensity (FI) per cell 24 h after treatment with DEX/DEX_RITC_-ILM as quantified by image analysis of CLSM images (*n =* 20 cells). **(c)** Flow cytometer measurement of cells 24 h after treatment with DEX/DEX_RITC_-ILM. **(d)** Representative phase contrast live cell images showing the interaction of filipodia with the DEX-ILM droplets (1000 nm) and subsequent uptake. Red arrow indicates DEX-ILM droplets being uptaken.

We further quantified the kinetics of DEX-ILM droplet uptake using CLSM image analysis (Figure **3**). Most DEX-ILM droplets are uptaken within 2 h, with the uptake peaking around 24 h and remaining stable for at least 96 h. The stable fluorescence intensity indicates that there is no detectable secretion of the droplets as we would otherwise expect a decrease in fluorescence intensity over four days. Specifically, the 96 h sample had a medium exchange 24 h prior to the measurement to ensure cell viability. If secretion were to occur, the medium exchange would have caused a measurable decrease in fluorescence intensity. However, since there is no significant difference between 96 h to 24, 36 or 72 h, the organelles are not secreted but remain inside the cells (Figure **3c**).

**Figure 3.**
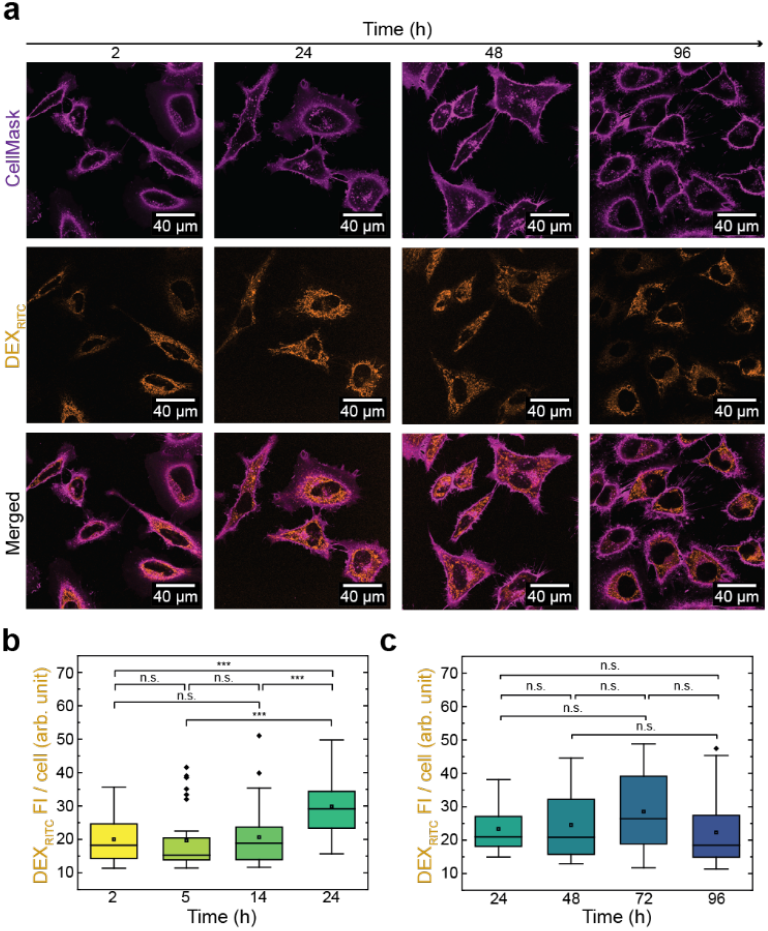
Time dependence of uptake of DEX/DEX_RITC_-ILM droplets (1000 nm). **(a)** CLSM images of HeLa cultures incubated for different periods after treatment, imaged with cell membrane stain (CellMask). **(b-c)** DEX_RITC_ FI per cell for samples incubated **(b)** < 24 h and **(c)** > 24h as quantified by image analysis of CLSM images (*n =* 30 cells). Graphs **b** and **c** show periods of continuous incubation; however, different microscope settings needed to be applied in both periods.

The cells remain viable, without adverse effects of the DEX-lipid droplets on cell morphology. Permeability assays with DAPI and flow cytometry (Figure **S2**) underscore high cell viability during 24 h (>90%; mostly above 96%), independent of the size of the DEX droplets or lipids used. This demonstrates that DEX-lipid droplets are overall well tolerated by the cells.

### Intracellular Stability of DEX Organelles and Orthogonal Organelles

One of the key questions for intracellular condensate-like organelles is whether they can undergo fusion inside the cytosol. To address this question, we added differently labeled DEX/DEX_RITC_-ILM and DEX/DEX_FITC_-ILM droplets (processed through 1000 nm membranes) to the same cells and evaluated colocalization of the fluorescent signals over time (see Experimental section for description of the statistical methods; Figures **4a**, **S3a**). The intensity scatter plot, correlating the intensity of each fluorescence channel for each pixel of a representative sample, does not show any correlation corresponding to colocalization that would be visible on the diagonal (Figures **4b**, **S3b**). This behavior is similar for 24 h and 96 h. We further measured the Pearson’s coefficient of the fluorescence distribution of both channels in single cells to quantify correlation. The Pearson’s coefficient remains < 0.5 up to 96 h, hence in a regime where correlation is absent (Figures **4c**, **S3c**). DEX-DOPC shows similar results (Figure **S4**). In summary, the DEX organelles do not fuse within the cell, highlighting their potential to build orthogonal organelle compartments. This property may be desirable for the implantation of parallel molecular systems into cells in the future.

**Figure 4.**
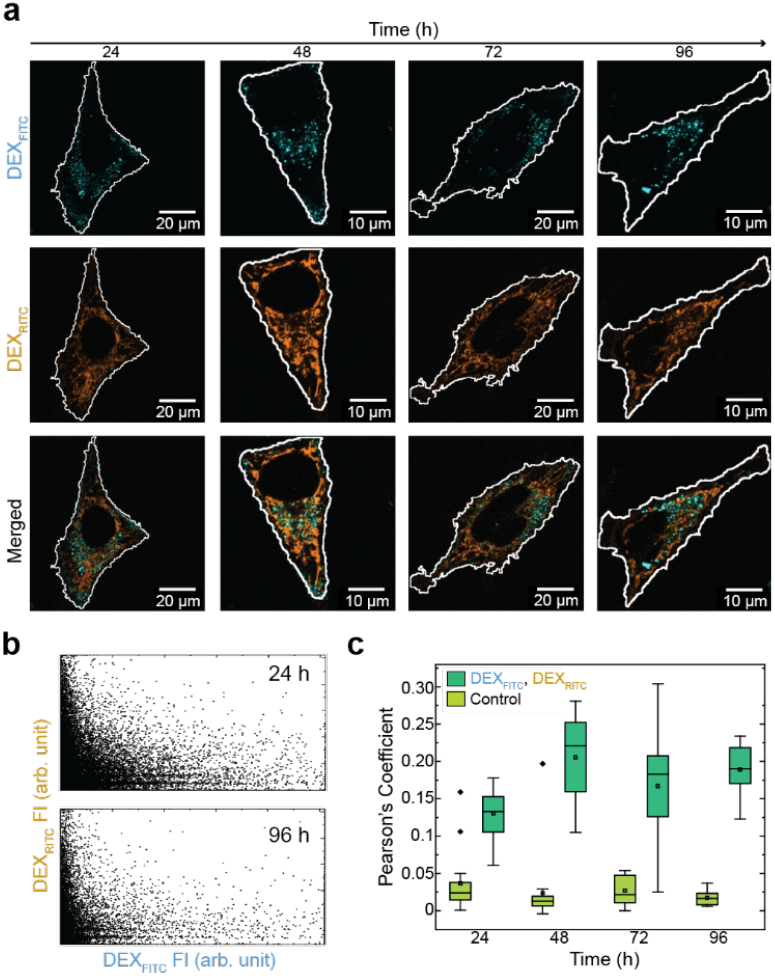
Organelle fusion is absent in cell. **(a)** Analysis of DEX/DEX_FITC_-ILM (1000 nm) and DEX/DEX_RITC_-ILM (1000 nm) droplet fusion inside cells by CLSM. Droplets containing different fluorescent tracers were mixed 1:1 v/v and given to HeLa cells and observed after incubation for 24, 48, 72 or 96 h. **(b)** Representative intensity scatter plot, showing fluorescence intensity per pixel of the RITC and FITC channels of a cell at 24 and 96 h. **(c)** Pearson’s correlation coefficient of the fluorescence intensity of the FITC and RITC channels for a population of cells (*n =* 16), control shows the same method applied to untreated cells (correlation of background fluorescence of the RITC and FITC channel).

### mRNA-Loaded DEX Organelles Enable Protein Translation

Dextran in DEX/PEG ATPS is known to spontaneously uptake large nucleic acids with high efficiency.^[43]^ We leveraged this property for the integration of eGFP-encoding mRNA into the ATPS dextran phase, that is subsequently incorporated into the DEX-lipid droplets (Figure **1**). By using a fluorescently labeled mRNA_AZ405-rUTP_ variant, we first observe that the mRNA remains inside the DEX organelles after cellular uptake (Figure **5a**). This is visible through the droplet-like shape of the cyan channel that visualizes the mRNA_AZ405-rUTP_ location.

**Figure 5.**
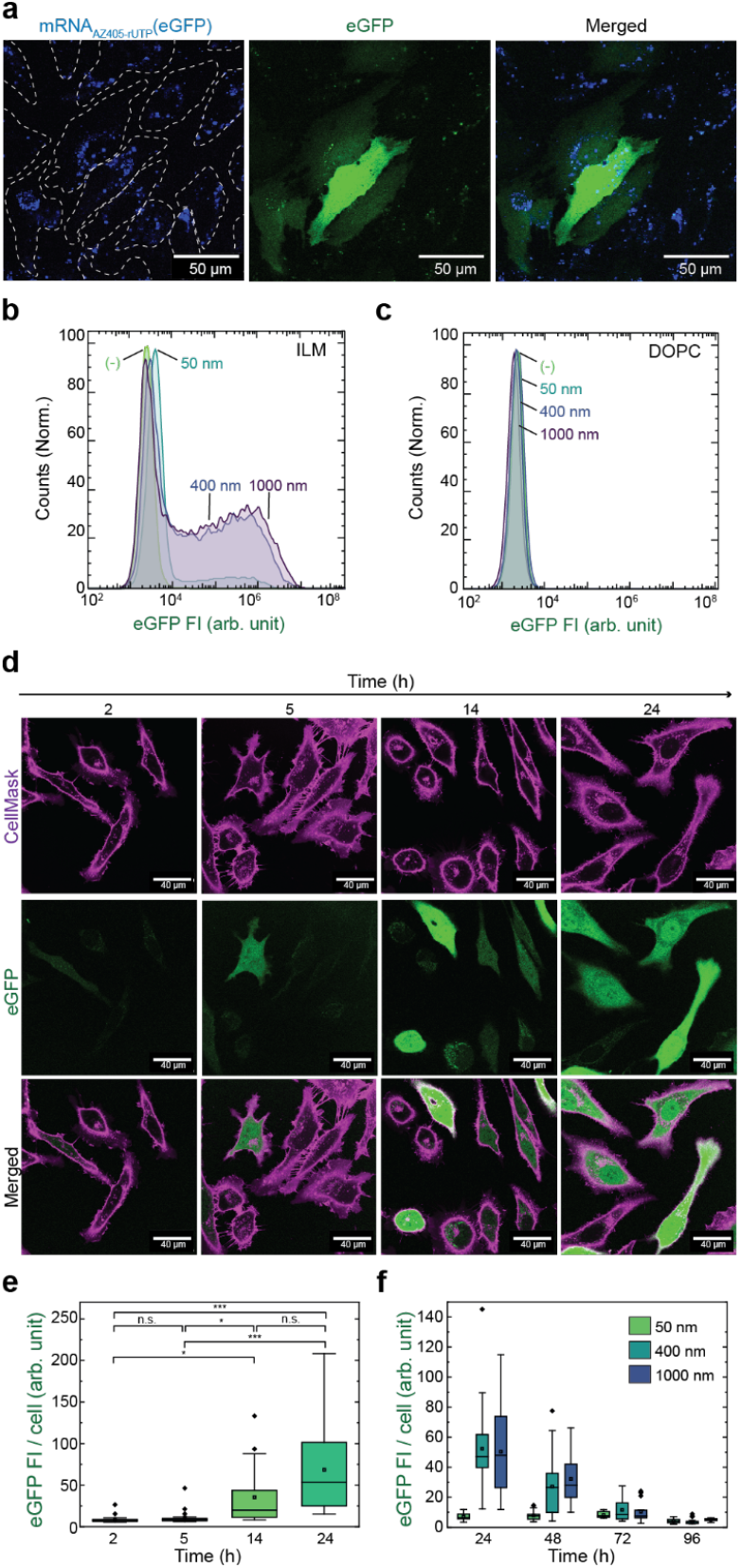
DEX-ILM Organelles have access to the translation machinery and produce eGFP. **(a)** CLSM images showing the localization of fluorescent mRNA delivered in DEX/mRNA_AZ405-rUTP_-ILM (1000 nm) and the resulting eGFP expression, dashed lines indicate the cell boundaries. **(b)** Flow cytometry of HeLa cells 24 h after treatment with DEX/mRNA-ILM showing the relation between eGFP levels and DEX-ILM droplet size. **(c)** Flow cytometry of cells treated with the DEX/mRNA-DOPC, showing absence of eGFP. **(d)** CLSM images of HeLa cells 24 h after treatment with DEX/mRNA-ILM (1000 nm), stained at the membrane with CellMask. **(e)** Distribution of eGFP fluorescence intensity per cell at incubations times ≤ 24 h with DEX/mRNA-ILM (1000 nm); quantification by image analysis of CLSM images (*n =* 30 cells). **(f)** Distribution of eGFP fluorescence intensity per cell at incubation times ≥ 24 h with DEX/mRNA-ILM according to extrusion size; quantification by image analysis of CLSM images (*n =* 30 cells). Graphs **e-f** show periods of continuous incubation, not continuous imaging of a single sample. Data within the same graph were prepared from cell cultures split from the same passage and treated at the same starting point.

Strikingly, the DEX/mRNA-ILM and DEX/mRNA_AZ405-rUTP_-ILM organelles allow for the translation towards eGFP, as found by flow cytometry and CLSM (Figures **5b,d,e**). A careful inspection of the CLSM images in Figure **5a** also highlights eGFP inside translationally active droplet, confirming the translation to take place inside the DEX-ILM organelles. In contrast, DEX/mRNA-DOPC do not lead to eGFP production (Figure **5c**). This confirms a fundamentally different fate of both organelles when entering cells. While both enter the cells, the ILM-based formulation that is optimized for endosomal escape indeed enables transition as organelles into the cytosol. It also shows that DEAE-DEX, added in trace amounts (∼1% w/w total DEX) to both formulations for better uptake of mRNA into the DEX droplets, is insufficient to induce endosomal escape for the DEX-DOPC droplets. The expression levels of protein and the non-toxicity observed for DEX-ILM droplets underscore that they can be possibly used for therapeutic mRNA transfection. The fact that the mRNA is translated but remains compartmentalized after uptake (Figure **5a**) is also relevant, since intracellular compartmentalization of mRNA in synthetic structures has been proposed to increase the efficiency of mRNA vaccines, and may contribute to its stability.^[44]^

We also analyzed the effect of droplet size on eGFP expression (Figure **5b**). Droplets prepared by extrusion through 400 nm and 1000 nm pores elicit the greatest eGFP expression, while those made with 50 nm pores have low expression. The decrease in eGFP levels of 50 nm DEX-ILM is not proportional to the decrease in uptake due to size. The uptake of 50 nm droplets is 49 % lower than for 1000 nm droplets (Figure **2b**), but eGFP expression is 92 % lower (Figure **5f**), suggesting that droplet size is not only relevant for overall uptake, but also for mRNA translation. This may relate to size-dependent effects on endosomal escape. For 400 nm and 1000 nm DEX-ILM droplets, approximately 30% of the cells show zero eGFP (Figure **5b**), indicating that cell cycle might also influence the mRNA translation.

The eGFP expression begins a few hours after treatment and can be seen on Supplementary Video **4**. It is pronounced after 5-14 h and peaks at 24 h (Figure **5e**). Decay of the protein fluorescence occurs similarly in cells that received 400 nm and 1000 nm sized droplets (Figures **5f**, **S5**).

### DEX organelles in key immune cell models

Immune cells, such as primary T-cells, are notoriously difficult to transfect. One reason is their membrane composition, low in heparan sulfate and proteoglycans, negatively charged molecules which enhance the interaction of the cell membrane with cationic lipoplexes.^[45]^ This is a challenge for immunotherapies such as CAR-T therapy.^[46]^ To understand the suitability of our DEX droplets for immune cell engineering, we investigated Jurkat, a T-cell model, and THP-1, a monocyte model. One strategy for increasing the transfection efficiency in Jurkat is their activation, which causes a widespread rearrangement of the endocytic machinery.^[47]^ Jurkat activation is often done by anti-CD3/CD28 antibodies.^[48]^ For THP-1 cells, a differentiation with Phorbol 12-myristate 13-acetate (PMA) causes the cells to become adherent, with a macrophage morphology.^[49]^

First, we tested the ability of the DEX-lipid droplets to be uptaken by the cells and form compartments. To this end, we treated the cells with DEX/DEX_RITC_-ILM (1000 nm) and measured the DEX_RITC_ fluorescence. The Jurkat and THP-1 cell lines quickly uptake the droplets. Like HeLa cells, a large fraction of the droplet is uptaken after 2 h (Fig. **6a,b**), with extensive uptake after 24 h. Cell viability remains acceptable at 78.9% 24 h after treatment (Figure **S6**), comparable to lipofection for these cell lines.^[50]^ Subsequently, we treated the cells with DEX/mRNA(mCherry)-ILM before and after activation or differentiation, respectively. We used mCherry instead of eGFP and doubled the mRNA concentration to achieve a better signal-to-noise ratio even at lower expression levels. Flow cytometry and CLSM clearly demonstrate a successful translation of the embedded mRNA in both Jurkat and THP-1 cells. Contrary to expectations, the pre-activation/differentiation of the cells has a negligible effect on mCherry translation (Fig. **6c,d**). Together, these results confirm that DEX-ILM droplets can serve as synthetic organelles in major immune cell models, expanding the scope for their use in future translational studies.

**Figure 6.**
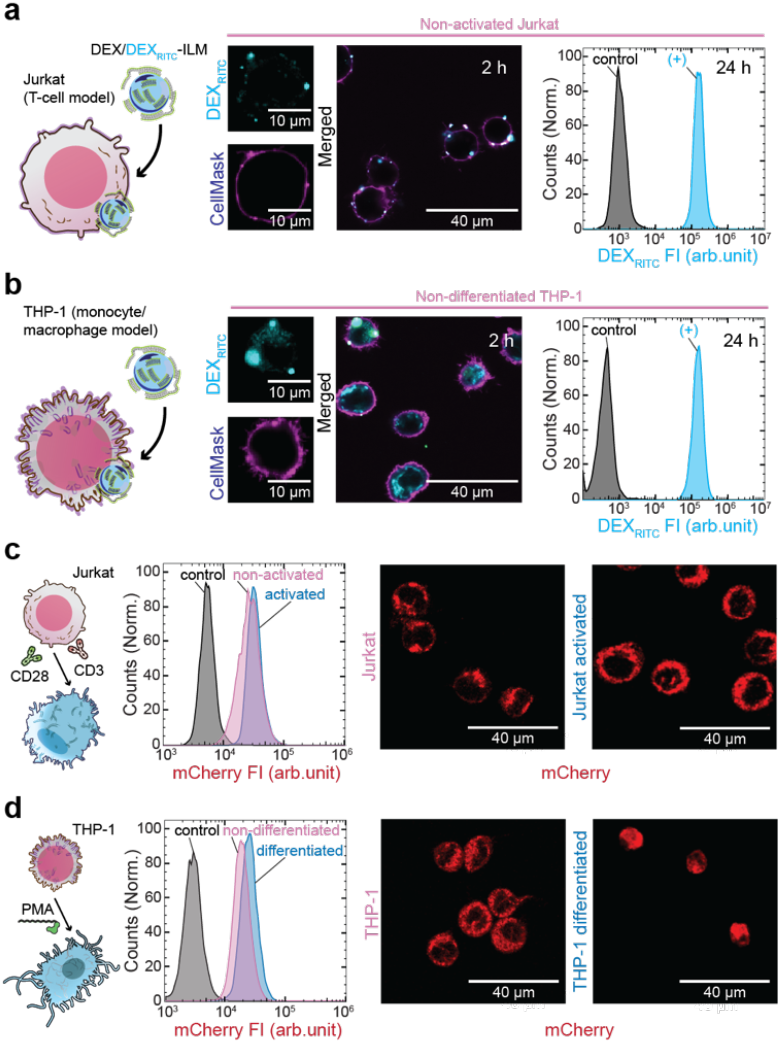
Uptake of DEX-lipid droplets and expression of reporter mRNA encoding mCherry in Jurkat and THP-1 immune cells. (**a-b)** Uptake of DEX/DEX_RITC_-ILM particles (1000 nm) by (a) Jurkat T-cells and (b) THP-1 monocytes. Center: Representative CSLM images show DEX_RITC_ localization (cyan) after 2 h, with membranes stained by CellMask Deep Red membrane stain (magenta). Right: Flow cytometer measurement of uptake after 24 h compared to untreated control. **(c-d)** Expression of mCherry in (c) Jurkat and (d) THP-1 cells 24 h after treatment with DEX/mRNA-ILM. Center: Flow cytometer measurement of mCherry fluorescence. Right: Representative CLSM images showing mCherry fluorescence (red). Jurkat: Non-activated and activated with anti-CD3 and anti-CD28 antibodies. THP-1: Non-differentiated and differentiated into macrophages with PMA. Negative controls as comparisons.

## Conclusion

Engineering the intracellular space is an emerging field of biomaterials research aimed at modifying cellular behavior. In parallel to endogenously produced condensates, we propose that there are advantages to producing components exogenously and introducing them into the cell, namely scalability, safety and flexibility in composition. We introduced DEX-lipid droplets prepared from a lipid-stabilized ATPS as a general and facile method to introduce designer condensates into cells. We showed that DEX-lipid droplets are uptaken by HeLa cells, are non-toxic and do not fuse within the cell, exhibiting the ability to form orthogonal compartments.

The droplets can be produced easily and exhibit self-assembling properties, such as encapsulation of an mRNA cargo. By adding ionizable lipids as droplet stabilizers, corresponding DEX-ILM droplets are capable of endosomal escape and transitioning into the cytosol. As a result, encapsulated mRNAs have access to the translation machinery and are translated while compartmentalized. We further demonstrated the uptake of the droplets and the transcription of the reporter mRNA in key immune cell models, pointing to potential applications in immunotherapies. The overlap of counterintuitive properties, such as avoidance of the endosome recycling, leads to novel research directions, such as investigating the molecular structure of DEX-lipid droplets, the uptake process, the role of filipodia, and the fate of transfected mRNA. We aim to clarify if other components of the cell interact with the droplet, such as ribosomes, which have affinity to DEX in a DEX-PEG ATPS.^[51]^ In the future, we believe dextran-lipid droplets can serve as a scaffold for the fabrication of more complex and functional cellular microreactors in other cell lines and *in vivo*.

## Supporting information

SupInfo

## Acknowledgements

M.M. acknowledges funding through the European Union’s Horizon 2022 research and innovation program under the Marie Skłodowska-Curie grant agreement DNAPC4ImunoMol-101111348. A.W. acknowledges a Max Planck Fellowship and the Gutenberg Research College and Deutsche Forschungsgemeinschaft (DFG, German Research Foundation) – 497845157 for funding the Zeiss Elyra 7. We acknowledge the IMB Flow Cytometry Core Facility for their support.

## Notes

### Competing Interest Statement

The authors have declared no competing interest.

