## Supplementary material for "Stable Synthetic Organelles from Aqueous Two-Phase Systems with Access to the Cell Translation Machinery": SupInfo

+Co-first authors

[a] L. Duttenhofer, Dr. M. Masukawa, Prof. Dr. A. Walther.

Life-Like Materials and Systems, Department of Chemistry

University of Mainz

Duesberweg 10-14, 55128 Mainz, Germany

[b] Dr. M. Masukawa, Prof. Dr. A. Walther

Max-Planck-Institute for Polymer Research

Ackermannweg 10, 55128 Mainz, Germany

#### Experimental Section

##### Materials

**Table S1.** Chemicals used in the fabrication of the DEX-lipid droplets.

| Abbreviation | Name | Supplier and product number |
|---|---|---|
| **DEX** | Dextran from Leuconostoc spp. MW 450,000-650,000 kDa | Sigma Aldrich, 31392-10G |
| **PEG** | Polyethylene glycol 6000 kDa | Sigma Aldrich, 528877-100GM |
| **FITC** | Fluorescein 5(6)-isothiocyanate, BioReagent | Fluka, 46950-1G-F |
| **RITC** | Rhodamine B isothiocyanate, BioReagent | Merck, R1755-100MG |
| **DEAE-DEX** | Diethylaminomethyl-Dextran 500,000 kDa | Sigma Aldrich, 93556-1G |
| **mRNA** | Messenger RNA encoding Enhanced Green Fluorescent Protein (Off-The-Shelf, dsRNA reduced, Cap1, 150 nt poly-A tail) | RiboPro, QS-U1-0003eGFP |
| **$mRNA_{AZ405\text{-}rUTP}$** | Fluorescently labelled Messenger RNA encoding Enhanced Green Fluorescent Protein (dsRNA reduced, Cap1, 25% AZ405-rUTP substitution, 150 nt poly-A tail) | RiboPro, O-CV1-LB1-AS-LN0-BW-QS-U1-0001 |
| **mRNA(mCherry)** | Messenger RNA encoding mCherry Fluorescent Protein (CleanCap™ mCherry mRNA, 5-methoxyuridine) | TriLink, L-7203-100 |
| **DOPC** | 1,2-Dioleoyl-sn-glycero-3-phosphocholine | Avanti Research, 850375P-25mg |
| **ALC-0315** | [(4-Hydroxybutyl)azanediyl]di(hexane-6,1-diyl) bis(2-hexyldecanoate) | Biomol, TGM-T9277-5mg |
| **Chol** | Cholesterol | Sigma Aldrich, C8667-5G |
| **DSPC** | 1,2-Distearoyl-sn-glycero-3-phosphocholine | Sigma Aldrich, P1138-1G |
| **$DOPE_{Atto488}$** | Atto488-1,2-Dioleoyl-sn-glycero-3-phosphoethanolamine | Atto-Tec, AD488-161 |
| **$PE_{Rhod}$** | 1,2-dioleoyl-sn-glycero-3-phosphoethanolamine-N-(lissamine rhodamine B sulfonyl) ammonium salt | Avanti Research, 810150C-1mg |

**Table S2.** Materials used in cell culture, microscopy and flow cytometer experiments.

| Abbreviation | Name | Supplier and product number |
|---|---|---|
| **HeLa** | HeLa (cervix carcinoma) cells | DSMZ, ACC57 |
| **Jurkat** | Jurkat (T-cell leukemia) cells | DSMZ, ACC282 |
| **THP-1** | THP-1 (acute monocytic leukemia) cells | DSMZ, ACC16 |
| **RNAse inhibitor** | Murine RNAse inhibitor | NEB, M0314S |
| **MEM** | Minimal Essential Media | Carl Roth, 9047.1 |
| **RPMI** | RPMI 1640 Medium (ATCC modification) | Thermo Fisher, A1049101 |
| **HEPES** | N-2-Hydroxyethylpiperazine-N'-2-ethane sulphonic acid 1M solution | Carl Roth, 9157.1 |
| **P/S** | Penicillin(10,000 units/mL), Streptomycin(10,000 µg/mL) solution | Fisher Scientific, 11548876 |
| **NEAA** | Non-essential aminoacids | Carl Roth, 9185.1 |
| **FBS** | Fetal Bovine Serum | Sigma, S0615-100ML |
| **BioTracker membrane** | BioTracker 490 Green Cytoplasmic Membrane Dye | Sigma Aldrich, SCT106 |
| **CellMask** | CellMask™ Deep Red Plasma Membrane Stain | Thermo Fisher, C10045 |
| **CellTrace** | CellTrace™ Violet | Thermo Fisher, C34557 |
| **Trypsin/EDTA** | ROTI®Cell Trypsin/EDTA solution (10x) | Carl Roth, 1Y19.1 |
| **Accutase** | StemPro™ Accutase™ Cell Dissociation Reagent | Thermo Fisher |
| **Anti-CD3** | Anti-Mouse CD3e monoclonal antibody (145-2C11) | Tonbo Biosciences, 70-0031-M001 |

| | | |
|---|---|---|
| **Anti-CD28** | CD28 Monoclonal Antibody (CD28.2), Functional Grade, eBioscience™, Invitrogen™ | Fisher scientific, 15278417 |
| **PMA** | Phorbol 12-myristate 13-acetate | Sigma Aldrich, P8139-1MG |

### Methods

#### Preparation of Fluorescent DEX

We prepared $DEX_{FITC}$ and $DEX_{RITC}$ according to de Belder and Granath.[4] Briefly, DEX (500 mg) was added to dimethyl sulfoxide (DMSO, 5 mL) and heated to 50 °C until completely dissolved. We added pyridine (60 µL), RITC or FITC (70 mg) and dibutyltin dilaurate (DBTL, 9 µL). We heated the mixture to 95 °C and stirred for 2 h. After cooling to RT, we precipitated and washed the product with cold EtOH three times, and freeze dried it. We estimated 100 glucose units per FITC for $DEX_{FITC}$ and 20 glucose units per RITC for $DEX_{RITC}$ by absorbance measurements.

#### Preparation of DEX-Lipid Droplets from Aqueous Two-Phase System

Table **S3** shows the components and final concentration of the base ATPS (1×)**.** Table **S4** specifies the composition of fluorescent dextran, lipids or mRNA in each experiment, in addition to the base ATPS. The only exception being Figure 1b, which has DEX at 1 mg/mL and $DEX_{RITC}$ at 0.1 mg/mL, to form large droplets and highlight surface accumulation of lipids. We start with an ATPS 2× concentrated, which produces micrometric DEX droplets. We used 1 mL of this emulsion to hydrate 200 µg of a lipid film. We prepared lipid films of either DOPC or an ionizable lipid mixture (ILM). The ILM consisted of ALC-0315:Chol:DSPC at a ratio 15:7:3 w/w/w. We prepared fluorescent variations of the lipid films by adding 1:1000 w/w of a fluorescent lipid. We used either $DOPE_{Atto488}$ or $PE_{Rhod}$. We prepared the lipid films by dissolving the lipids in chloroform in a round-bottomed glass tube (12 x 55 mm), then carefully drying them with a nitrogen stream while rotating the tube. We placed the films in vacuum for 2 h for complete evaporation of the solvent and stored them at -20°C until further use. Before using it, we brought the films to RT, poured the ATPS (1 mL) onto the films and vortexed (3000 rpm, 1 min, hand warmed). We then diluted the lipid-stabilized ATPS with cell culture medium at a ratio 1:1 v/v and added 600 units/mL of murine RNAse inhibitor. The medium consisted of MEM, 1% (v/v) HEPES, 1% (v/v) P/S, 1% (v/v) NEAA for HeLa cells, and RPMI, 10% FBS for Jurkat and THP-1. We performed the extrusion with a mini-extruder (Avanti Research 610000-1EA) 11 times, using polycarbonate membranes with pore sizes of 50 nm, 400 nm or 1000 nm. After extrusion, the droplets were once again diluted in medium with serum 1:1 v/v.

**Table S3.** Composition of the ATPS used as a base for the fabrication of DEX-lipid droplets.

| Component | Concentration |
|---|---|
| **PEG** | 100 mg/mL |
| **DEX** | 71 µg/mL |
| **DEAE-DEX** | 0.8 µg/mL |
| **(if added) $DEX_{RITC}$ or $DEX_{FITC}$** | 8 µg/mL |
| **(if added) mRNA or $mRNA_{AZ405-rUTP}$** | 8 µg/mL |
| **(if added) mRNA(mCherry)** | 16 µg/mL |
| **(if RNA added) RNAse inhibitor** | 600 U/mL |

#### Dynamic Light Scattering (DLS)

We diluted the extruded DEX-ILM droplets 1:1 v/v in PBS and measured their size distribution using a Malvern Zetasizer Nano ZS.

#### HeLa Cell Culture

We cultured HeLa cells in MEM supplemented with 1% v/v HEPES, 1% v/v P/S, 1% v/v NEAA and 10% v/v FBS at 37 °C in 5% $CO_2$ humidified air. We used 8-well slides with a seeding density of $2\times10^3$ cells per 300 µL, performing the experiments typically 24 h after seeding. We washed the cells with PBS and added the medium containing the DEX-lipid droplets. Composition of specific experiments are listed in Table **S4**. For experiments spanning 96 h, we performed one medium change 24 h prior to the measurements. For experiments < 96 h, no medium change was necessary. In microscopy experiments, we stained the cells with either BioTracker Membrane, CellMask or CellTrace. The membrane stains were used according to the manufacturer instructions, which recommend adding stains up to two hours prior to observations. Additionally, we used CellTrace, a typical stain for flow cytometry, as a microscopy stain at one-tenth of the recommended concentration.

#### Jurkat and THP-1 Cell Culture

We cultured Jurkat and THP-1 cells in RPMI supplemented with 10% v/v FBS at 37°C and 50 μM 2-mercaptoethanol (added fresh at every passage) in 5% CO2 humidified air. We cultured the cells in 5 ml suspension cell tubes with a seeding density of $1×10^5$ cells in 1 mL. When activating Jurkat cells, we coated the tube with 1 μg/mL anti-CD3 and 1 μg/mL anti-CD28 antibodies in cell media for 30 minutes prior to the addition of the cells. For differentiation of THP-1 cells, we added 100 ng/mL PMA. DEX-lipid droplets were added 24 h after seeding/stimulus. Prior to treatment with DEX-lipid droplets, the cells were centrifuged at 400 x g for 5 min, the medium was removed, and the DEX-lipid droplets were added. Composition of specific experiments are listed in Table **S4**. For the microscopy experiments, we stained the cells with CellMask at the concentration recommended by the manufacturer 30 minutes before imaging.

**Table S4.** Composition of the droplets used in each figure.

| Figure | $DEX_{FITC}$ | $DEX_{RITC}$ | $DOPE_{Atto488}$ | $PE_{Rhod}$ | mRNA(eGFP) | $mRNA_{AZ405-rUTP}$ | mRNA(mCherry) |
|---|---|---|---|---|---|---|---|
| **1b** | | + | + | | | + | |
| **1c** | | + | + | | | + | |
| **1d** | | | | | | | |
| **2a-c** | | + | | | | | |
| **2d** | | | | | | | |
| **3a-c** | | + | | | | | |
| **4a-c** | + | + | | | | | |
| **5a** | | | | | | + | |
| **5b-f** | | | | | + | | |
| **6a-b** | | + | | | | | |
| **6c-d** | | | | | | | + |
| **S1a-b** | | | | + | | | |
| **S1d** | | + | | | | | |
| **S2** | | | | | | | |
| **S3** | + | + | | | | | |
| **S4** | + | + | | | | | |
| **S5** | | | | | + | | |
| **S6** | | + | | | | | |

#### Flow Cytometry

We performed flow cytometry experiments 24 h after the addition of the DEX-lipid droplets. For HeLa cells, we removed the medium mixture from the cells, and washed them with 1 mL of PBS. We trypsinized the cells for 10 min with Trypsin/EDTA in PBS. We added 1 mL of the cell culture medium to quench the trypsinization and centrifuged the cells at ~750×g. We then removed the supernatant and suspended the cells in flow cytometer buffer (400 μL PBS, 2 mM EDTA, 2% (v/v) FBS). Jurkat and THP-1 cells were harvested directly from the culture tube for flow cytometer measurements, except for differentiated THP-1 cells, which were attached. For THP-1 cells, we removed the media, added 200 μL of Accutase for 3 min and added 800 μL of media. To examine cell viability, we added DAPI 0.1 μg/mL as a permeability marker and incubated for 30 min before measurement using an Agilent NovoCyte Quanteon. (Agilent, NovoExpress v.1.6.0) with 4 excitation lasers (405 nm, 488 nm, 561 nm and 640 nm). The raw data was processed and plotted using FlowJo 10.10.0.

#### Microscopy

We performed Confocal Laser Scanning Microscopy (CLSM) with a Leica Stellaris 5 microscope (LasX v4.3.0.24308) with four laser lines (405 nm, 488 nm, 561 nm and 638 nm). For fluorescence live cell imaging and phase contrast live cell imaging we used a Zeiss Elyra 7 equipped with onstage incubator. Leica Stellaris 5 microscope (LasX v4.3.0.24308) with four laser lines and three HyD S detectors using plan-apochromat objectives (63×, 1.40 numerical aperture, oil immersion). Elyra 7 Imaging System equipped with four laser lines (405 nm, 488 nm, 561 nm, and 642 nm) using alpha Plan-Apochromat 63x, N.A. 1.46.oil immersion, pco.edge 4.2 CLHS water-cooled sCMOS cameras. We used ImageJ[5] v1.54f and OriginPro 2023b for image analysis.

#### Droplet Fusion Image Analysis

We simultaneously incubated the cells with a mixture of two dextran-lipid droplets, one DEX/$DEX_{RITC}$-ILM and the other DEX/$DEX_{FITC}$-ILM. We used CLSM images to measure the correlation of $DEX_{RITC}$ and $DEX_{FITC}$ fluorescence. We used the Pearson's coefficient (PC) to quantify correlation and possible overlap between the droplets, using single cells as regions of interest (ROIs). The PC expresses the pixel-by-pixel covariance in the intensity of two channels and is given by the following equation[6]

$$PC = \frac{\sum_i (A_i - a) \times (B_i - b)}{\sqrt{\sum_i (A_i - a)^2 \times \sum_i (B_i - b)^2}}$$

where $Ai$ and $Bi$ are the intensity values of pixel $i$ in channel $A$ and $B$, respectively, while $a$ and $b$ represent the average intensity of the entire channel $A$ and $B$ of a given ROI. The PC takes values between -1, which indicates a perfect anti-correlation, and +1, which indicates a perfect correlation of both channels. Values between -0.5 and +0.5 do not allow any conclusions about colocalization to be drawn.

We performed Bonferroni-corrected post-hoc tests for the boxplots of PC to analyze the statistical significance within an ensemble of more than two data sets.[8] Therefore, we determined the *p*-values for each combination of data set pair by the *t*-test assuming equal variances. We then obtained the Bonferroni-corrected significance values by dividing the standard significance values by the total number of comparisons. Finally, the comparison of the *p*-values with the Bonferroni-corrected significance values yielded the respective significance level shown in the boxplots. Table **S5** shows the corresponding *p*-values, the Bonferroni-corrected *p*-values and the significance.

**Supplementary Table 5.** Correspondence between the *p*-values, Bonferroni-corrected significance values and significance levels applied to boxplot annotations.

| Standard *p*-value limits | Bonferroni-corrected *p*-value limits | Significance level |
|---|---|---|
| **>0.05** | >0.008333 | Not significant (n.s.) |
| **<0.05** | <0.008333 | * |
| **<0.01** | <0.001667 | ** |
| **<0.001** | <0.000167 | *** |

#### Supplementary Figures

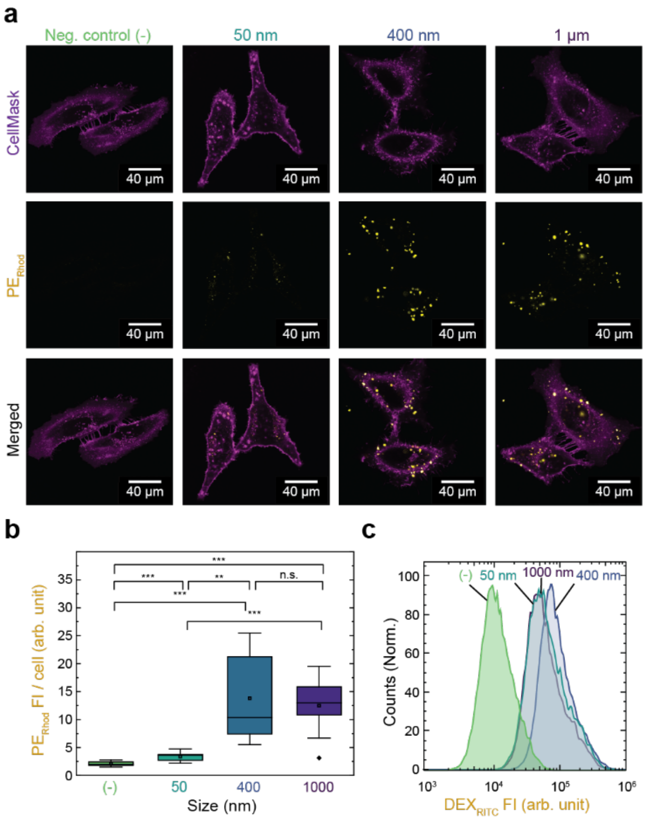


**Figure S1.** Uptake of DEX-DOPC droplets by HeLa cells according to extrusion size. **(a)** CLSM images of cells 24 h after treatment with DEX-DOPC/$PE_{Rhod}$ and cell membrane stain CellMask. Culture medium added as negative control. **(b)** Distribution of $PE_{Rhod}$ fluorescence intensity (FI) per cell 24 h after treatment with DEX-DOPC/$PE_{Rhod}$, quantification by image analysis of CLSM images ($n$ = 10 cells). **(c)** Flow cytometer measurement of cells 24 h after transfection with DEX/$DEX_{RITC}$-DOPC.

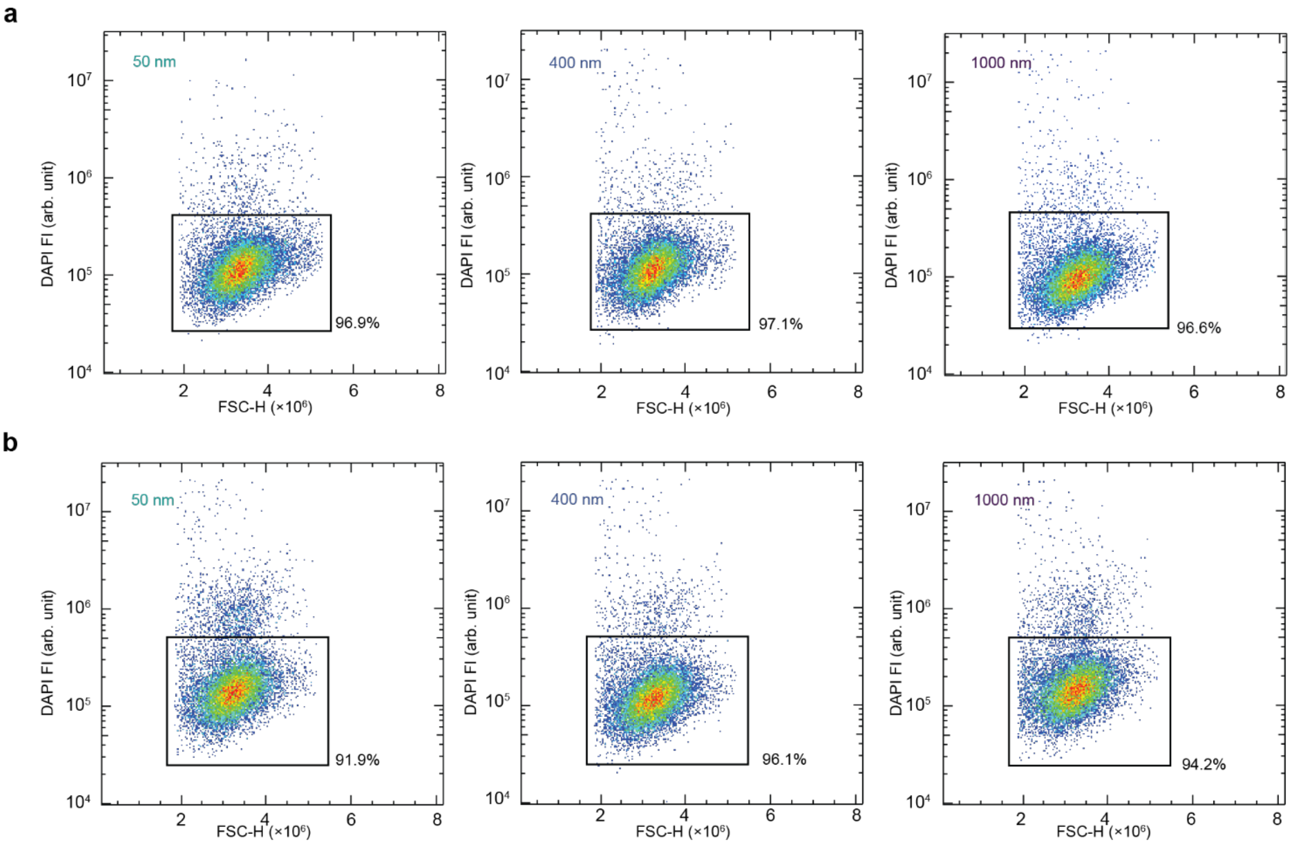


**Figure S2.** Density plot of flow cytometer viability assay of HeLa cells stained with DAPI, 24h after treatment with DEX-lipid droplets. Data points represent the maximum intensity of the forward scattering signal (FSC-H) relative to the DAPI fluorescence signal of individual cells. The annotations indicate the size of the DEX-lipid droplets used, and the gates show the percentage of live cells (DAPI negative). Droplets in the medium that were not taken up are removed during washing, and do not appear in the flow cytometer data. **(a)** Cells treated with DEX-ILM. **(b)** Cells treated with DEX-DOPC.

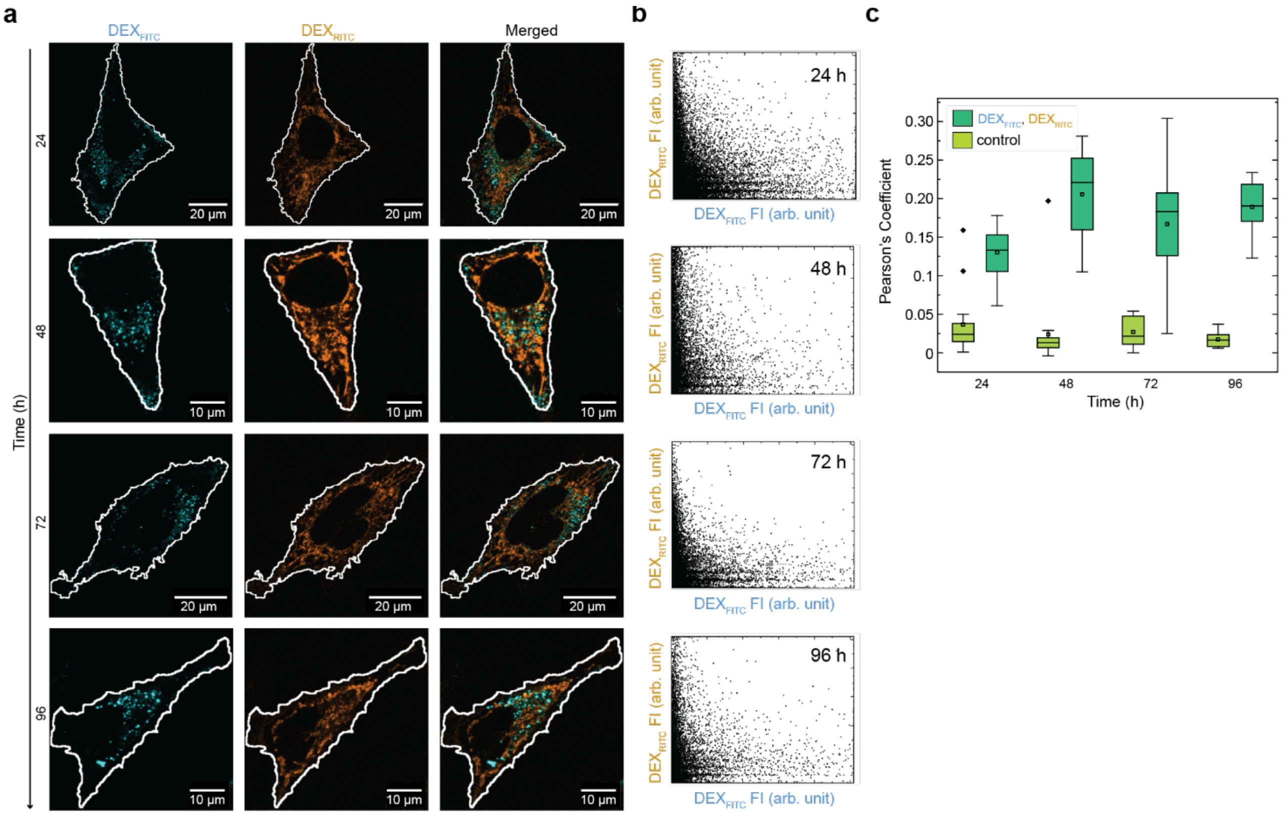


**Figure S3.** Droplet fusion assay, cells treated simultaneously with DEX/DEX$_{FITC}$-ILM (1000 nm) and DEX/DEX$_{RITC}$-ILM (1000 nm). Data partially reproduced in Fig. 4. **(a)** Representative CLSM images after different incubation intervals (24, 48, 72, 96 h). **(b)** Representative intensity scatter plots of the pixels in a single cell. *x*-values correspond to pixel intensity in the FITC channel, and *y*-values correspond to pixel intensity in the RITC channel. **(c)** Pearson's coefficient (correlation quantification) of the FITC, RITC channels at different time points obtained by image analysis, where the control corresponds to the untreated cells (correlation between the background noise of the channels), $n$ = 16 cells.

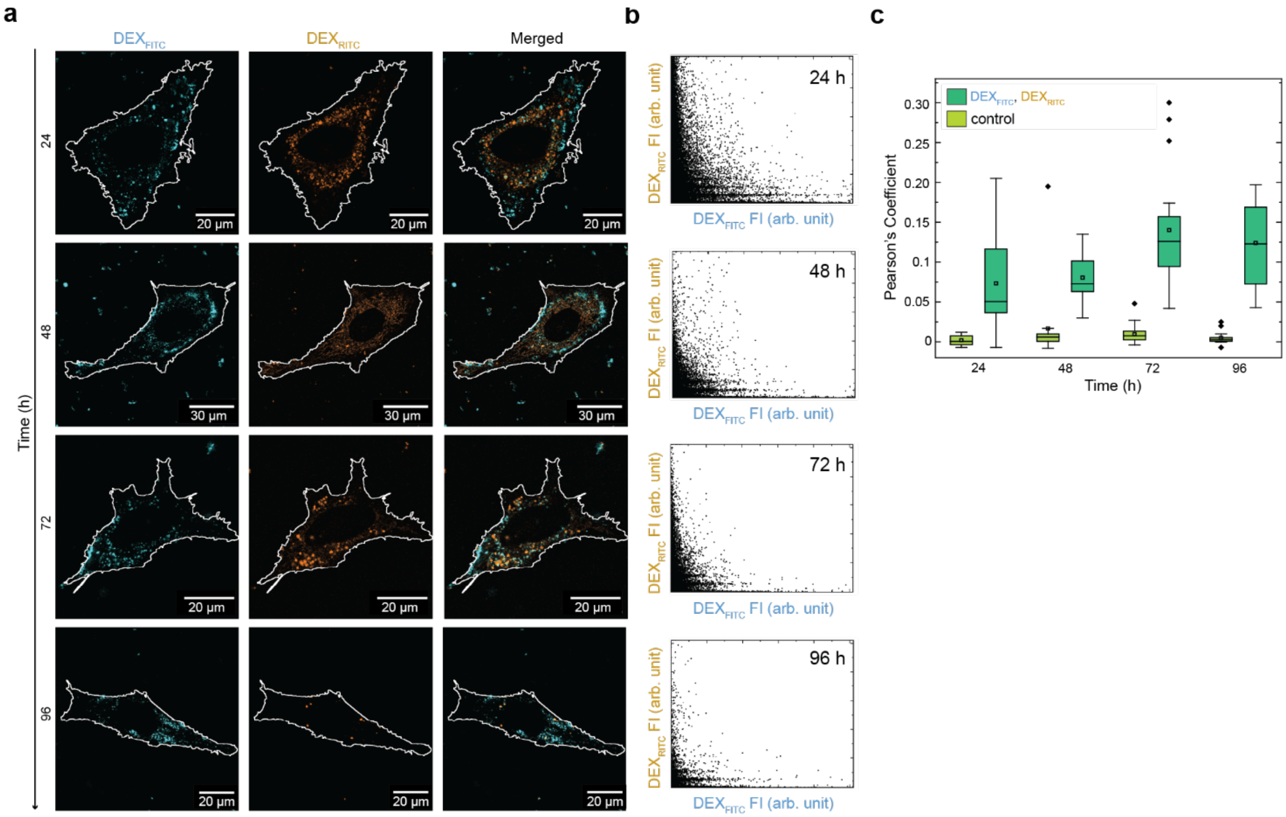


**Figure S4.** Droplet fusion assay, cells treated simultaneously with DEX/$DEX_{FITC}$-DOPC (1000 nm) and DEX/$DEX_{RITC}$-DOPC (1000 nm). **(a)** Representative CLSM images after different incubation times (24, 48, 72, 96 h). **(b)** Representative intensity scatter plots of the pixels in a single cell. *x*-values correspond to pixel intensity in the FITC channel, and *y*-values correspond to pixel intensity in the RITC channel. **(c)** Pearson's coefficient (correlation quantification) of the FITC, RITC channels at different time points, where the control corresponds to the untreated cells (correlation between the background noise of the channels), $n$ = 16 cells.

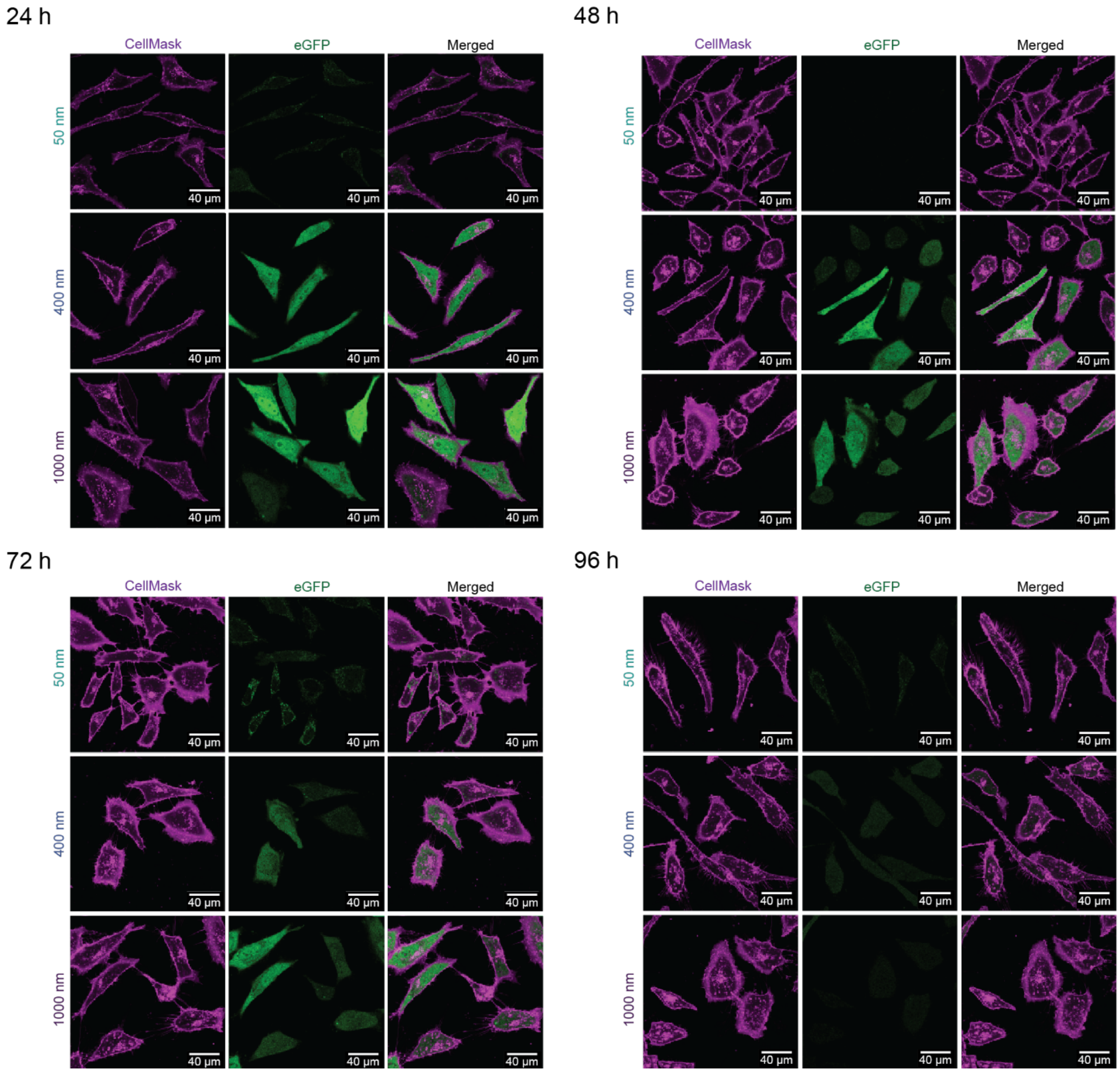


**Figure S5.** Representative CLSM images of cells treated with DEX/mRNA-ILM droplets extruded with 40, 50 or 1000 nm pore size membranes, and incubated for 24, 48, 72 or 96 h, with membrane stain CellMask.

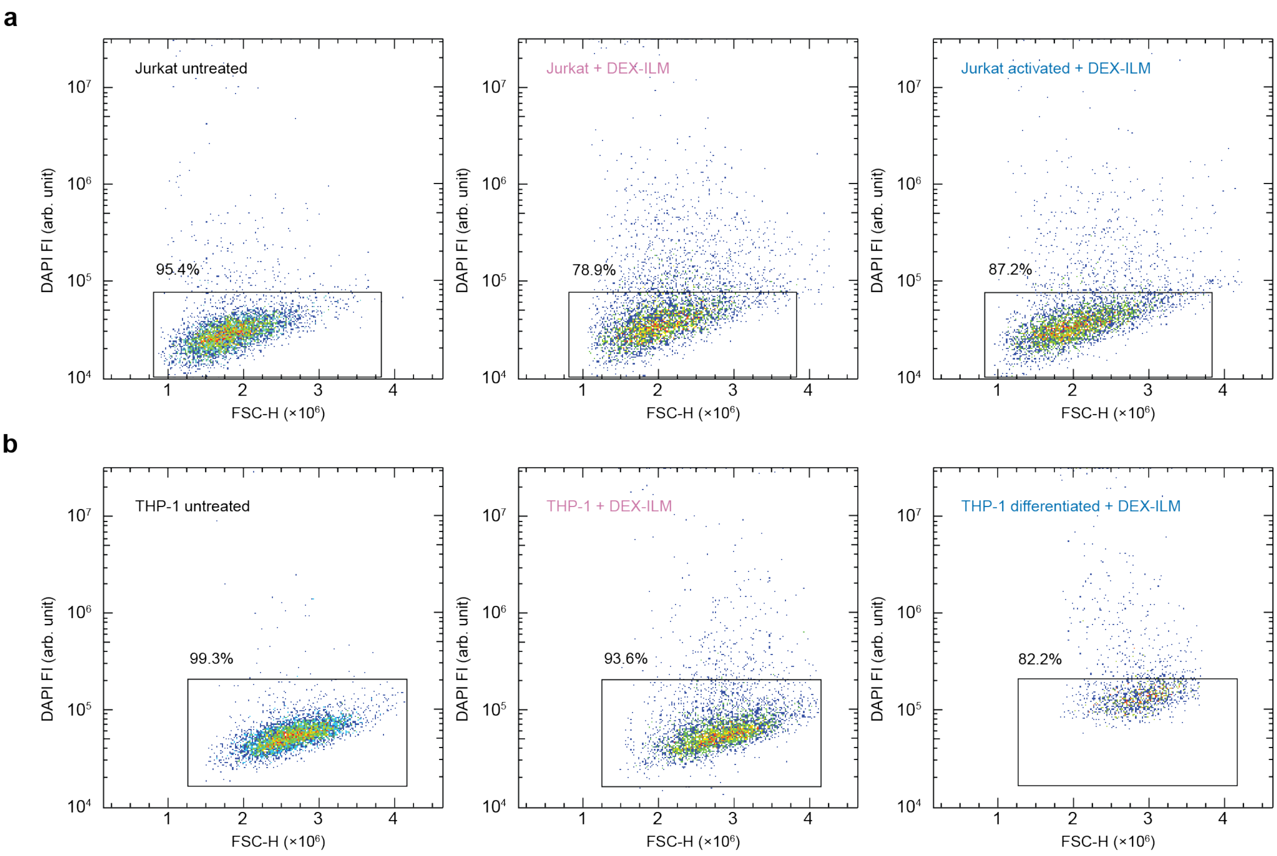


**Figure S6.** Density plot of flow cytometer viability assay with DAPI, 24h after treatment with DEX-ILM droplets. Data points represent the maximum intensity of the forward scattering signal (FSC-H) relative to the DAPI fluorescence signal of individual cells. Gates indicate the percentage of DAPI negative cells (viable) **(a)** Jurkat cells. **(b)** THP-1 cells. Annotations indicate the cell type and stimuli. Untreated refers to cells that did not receive the DEX-lipid droplet or stimuli. Activated Jurkat cells stimulated with anti-CD3/CD28 antibodies, differentiated THP-1 stimulated with PMA 48 h prior to measurement.

#### Supplementary Videos

**Supplementary Video 1. a.** Fluorescence live cell imaging of cells treated with DEX-DOPC/$PE_{Rhod}$ (1000 nm). Magenta channel: BioTracker 490 Green Cytoplasmic Membrane Dye stain. Yellow channel: $PE_{Rhod}$. **b.** Magnification of **a.**

**Supplementary Video 2.** Fluorescence live cell imaging of cells treated with DEX/ $DEX_{RITC}$-ILM (1000 nm). Magenta channel: CellTrace Violet. Orange channel: $DEX_{RITC}$.

**Supplementary Video 3.** Phase contrast live cell imaging of cells treated with DEX-ILM (1000 nm).

**Supplementary Video 4.** Fluorescence live cell imaging of the initial hours of cells treated with DEX/$DEX_{RITC}$/mRNA -ILM (1000 nm). Orange channel: $DEX_{RITC}$. Green channel: eGFP.
